# Reduced Cognitive Performance in Aged Rats Correlates with Increased Excitation/Inhibition Ratio in the Dentate Gyrus in Response to Lateral Entorhinal Input

**DOI:** 10.1101/637439

**Authors:** Trinh Tran, Michelle Bridi, Ming Teng Koh, Michela Gallagher, Alfredo Kirkwood

## Abstract

Aging often impairs cognitive functions associated with the medial temporal lobe (MTL). Anatomical studies identified the layer II pyramidal cells of the lateral entorhinal cortex (LEC) as one of the most vulnerable elements within the MTL. These cells provide a major excitatory input to the dentate gyrus hippocampal subfield by synapsing onto granule cells and onto local inhibitory interneurons, and a fraction of these contacts are lost in aged individuals with impaired learning. Using optogenetics we evaluated the functional status of the remaining inputs in an outbred rat model of aging that distinguishes between learning impaired and learning unimpaired individuals. We found that aging affects the pre- and postsynaptic strength of the LEC inputs onto granule cells. However, the magnitude these changes was similar in impaired and un-impaired rats. In contrast, the recruitment of inhibition by LEC activation was selectively reduced in the aged impaired subjects. These findings are consistent with the notion that the preservation of an adequate balance of excitation and inhibition is crucial for maintain proficient memory performance during aging.

## INTRODUCTION

Aging often has a negative impact on cognitive abilities, particularly on learning and the encoding of new memories. It was early recognized that these impairments occur in the absence of significant neuronal loss, but are more likely due to alterations in the strength, connectivity and plasticity of circuits within the medial temporal lobe (MTL) (Rapp and Gallagher, 1996), which comprises the hippocampal formation and medial temporal cortex (entorhinal, perirhinal and parahippocampal regions). Within the MTL memory system proficient episodic memory critically depends on the computational properties of the dentate gyrus that receives highly processed input from layer II neurons of the entorhinal cortex. This pathway provides excitatory input to both excitatory granule cells and inhibitory interneurons in the dentate gyrus. Information processing in the dentate gyrus performs pattern separation, referring to the highly distinctive encoding of input in a sparse network of granule cells (Leutgeb et al., 2007; McHugh et al., 2007; Neunuebel and Knierim, 2014). Such encoding is crucial to minimize interference between representations of similar but not identical experiences in episodic memory.

Recent studies have revealed that in elderly humans, memory loss and mild cognitive impairment correlates with reduced structural connectivity between the entorhinal cortex and the DG (Scheff et al. 2006; Yassa et al., 2010a), a diminished capacity for pattern separation in tests of memory performance, and hyperactivity in the dentate/CA3 subfields of the hippocampus (for review see Leal and Yassa, 2015). Indeed, hyperactivity appears to be a dysfunctional condition common to cognitive impairment in aged rodents (Wilson et al., 2005), mouse models of AD (Palop et al., 2007) and elderly humans diagnosed with mild cognitive impairment (Yassa et al., 2011; Bakker et al. 2012) potentially contributing to loss of pattern separation in the DG.

Although much evidence has indicated a reduction in the Layer II input to the DG in aging based on studies in both laboratory animals and humans (Bondareff & Geinisman 1976; Rapp et al. 1999; Smith et al. 2000; Yassa et al. 2010a), few studies have examined the functional properties of the remaining circuitry in aging (but see Foster et al 1991). Here we focus on the input to the DG from the lateral EC (LEC) that innervates granule cell dendrites in the outer molecular layer of the DG and provides feedforward inhibition via interneurons in the DG. Anatomical and structural studies in aged rats and humans indicate that the layer II pyramidal cells in the LEC are specifically affected in memory impaired individuals (Stranahan et al., 2010;) Stranahan et al., 2011; and age-related effects on this input are augmented in both prodromal and early dementia phases of late onset AD when spreading of tau pathology initially localized in the lateral EC occurs (Scheff et al. 2006; Khan et al., 2014; Tward et al. 2018; Kulason et al. 2019).

Motivated by the considerations above we set out to evaluate how age affects the strength of synaptic excitation and synaptic inhibition in response to LEC→GC inputs. The study was performed in a model of behaviorally characterized outbred aged rats that distinguish between impaired and unimpaired individuals (Gallagher et al. 1993). The selective activation of LEC inputs was achieved with optogenetics. We found that aging affected both pre and postsynaptic function of the direct LEC excitatory inputs onto granule cells, but to a similar extent in behaviorally impaired and unimpaired aged individuals. In contrast, the recruitment of disynaptic inhibition by LEC activation was selectively reduced in the aged impaired subjects. This finding is consistent with the notion that the preservation of an adequate balance of excitation and inhibition to control sparse encoding in DG is crucial for maintain proficient memory performance during aging.

## METHODS

### Behavioral assessment

Male Long-Evans outbred rats obtained pathogen-free from Charles River Laboratories (Raleigh, N.C.) were 6 month (young) or 24 month (aged) of age at the time of behavioral characterization for spatial learning in a water maze (1.83 m diameter, opaque water at 27°C). During an eight-day period, in sessions consisting of three trials a day with a 60 s inter-trial interval, rats were trained to locate a camouflaged platform that remained in the same location 2 cm below the water surface. During a training trial, the rat was placed in the water at the perimeter of the pool and allowed 90 s to locate the escape platform. If at 90 s the rat failed to escape on a trial, it was placed onto the platform and allowed to remain there for 30 s. The position of entry for the animal was varied at each trial. Every sixth trial consisted of a free swim (“probe trial”), which served to assess the development of a spatially localized search for the escape platform. During probe trials the rat was allowed to swim a total of 30 s with the escape platform retracted to the bottom of the pool. After 30 s, the platform was raised so that the rat could complete escape on the trial. A “behavioral index”, which was generated from the proximity of the rat to the escape platform during probe trial performance, was used in correlational analysis with the neurobiological data. This index is the sum of weighted proximity scores measured during probe trials; low scores reflect search near the escape platform whereas high scores reflect search farther away from the target. Thus, the “behavioral index” provides a measure that is based on search accuracy independent of escape velocity (Gallagher, 1993). “Search error” during training trials refers to the deviation from a direct path to the platform and provided an additional measure for behavioral analysis (Gallagher et al. 1993), Cue training (visible escape platform; 6 trials) occurred on the last day of training to test for sensorimotor and motivational factors independent of spatial learning. Rats that failed to meet a cue criterion of locating the visible platform within an averaged of 20 s over six trials were excluded from the experiments.

### Viral Transfection

Rats were anesthetized with isoflurane mixed with O2 and transcranially injected bilaterally with 0.5 µl adeno-associated virus containing channelrhodopsin-2 and yellow fluorescence protein as a marker (AAV2/9.hSynapsin.hChR2(H134R)-EYFP.WPRE.hGH, Addgene26973, Penn Vector Core) into the lateral entorhinal cortex at coordinates: (1) Bregma −5.2 mm, Lateral ±7.0 mm, Depth −9.0, and (2) Bregma −6.3 mm, Lateral ±6.0 mm, Depth −8.0 mm. Rats recovered on a heated surface and were returned to the animal colony, where they remained for 6-7 weeks to allow optimal ChR2 expression.

### Brain slices

All electrophysiological studies were performed by experimenters who were blinded to the behavioral score of the subject. Behaviorally characterized young (6 months) and aged (24 months) rats were deeply anesthetized with isoflurane followed by urethane anesthesia (1 g/kg, i.p.), and perfused transcardially with cold dissection buffer (75 ml at 25 ml/min) containing the following (in mM): 92 N-methyl-D-glucamine (NMDG), 2.5 KCl, 1.25 NaH_2_PO_4_, 30 NaHCO_3_, 20 HEPES, 25 glucose, 2 thiourea, 5 Na-ascorbate, 3 Na-pyruvate, 0.5 CaCl_2_ and 10 MgSO_4_, pH adjusted to 7.4 with HCl. Rats were then decapitated and the brains were removed quickly. Coronal hippocampal slices (300μM) were prepared as described (Boric et al., 2008) in ice-cold dissection buffer bubbled with a mixture of 5% CO_2_ and 95% O_2_. Slices were incubated for 15 minutes at 32°C in dissection buffer then allowed to recover for 1 hr at room temperature in artificial cerebrospinal fluid (ACSF) containing the following (mM): 124 NaCl, 5 KCl, 1.25 NaH_2_PO_4_, 26 NaHCO_3_, 10 dextrose, 1.5 MgCl_2_, and 2.5 CaCl_2_, bubbled with 95% O_2_ / 5% CO_2_. All procedures were approved by the Institutional Animal Care and Use Committee at Johns Hopkins University.

### Visualized whole-cell voltage-clamp recordings

All recordings were done in a submerged recording chamber superfused with ACSF (30 ± 0.5°C, 2 ml/min) in the presence of 100 μM APV. In the experiments reported in figure 2, 5 mM SrCl_2_ and 10 µM bicuculine methiodide (BMI) were also added. Visualized whole-cell voltage-clamp recordings in DG granule cells with glass pipettes (4–6MΩ) filled with intracellular solution containing the following (in mM): 120 CsCl, 8 NaCl, 10 HEPES, 2 EGTA, 5 QX-314, 0.5 Na2GTP, 4 MgATP, and 10 Na2-phosphocreatine, pH adjusted to 7.25 with KOH, 280–290 mOsm. Membrane currents were recorded at −70 mV in the presence 100 µM 2-amino-5-phosphonovaleric acid (APV). Cells with series resistance > 25 MΩ, input resistance < 100 MΩ, and/or RMS noise > 4pA were excluded from analysis. Membrane resistance and series resistance were monitored with 100-msec negative voltage commands (−4 mV) delivered before the stimulation. Cells were also excluded if series resistance changed > 15% over the course of the experiment. Data were filtered at 2 kHz and digitized at 10 kHz using Igor Pro (WaveMetrics Inc., Lake Oswego, OR). All drugs were purchased from Sigma or R&D (Tocris).

**Figure 1.**
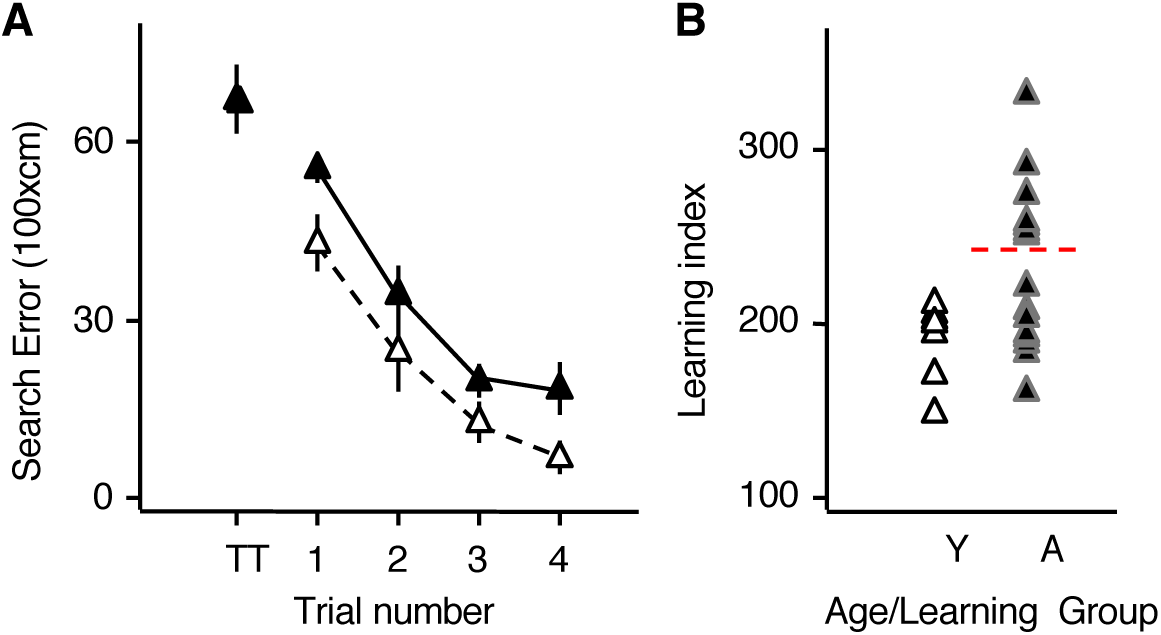
Behavioral characterization of young and aged rats in the spatial version of the Morris water maze. **A**) Cumulative search error measure of learning in five trial blocks during water maze training. This measure reflects the distance of the rat from the escape platform throughout its search, with higher numbers indicating worse performance. On the initial training trial (TT) there was no difference in the performance of young and aged rats. Statistical analysis described in the text indicated that the aged rats overall performed more poorly than young over the course of training. Data points represent the average for blocks of five training trials ± SEM. **B)** A learning index measure for each rat was derived from proximity of the rat’s search during probe trials interpolated throughout training as described in Gallagher et. al., 1993, with lower scores indicating more accurate performance. As a group aged rats exhibited significant impairment in accuracy and a larger variability of individual outcomes. Nearly half of the aged rats performed more poorly than young rats (designated aged impaired), while the rest performed within the range of younger adults (designated aged unimpaired).

**Figure 2.**
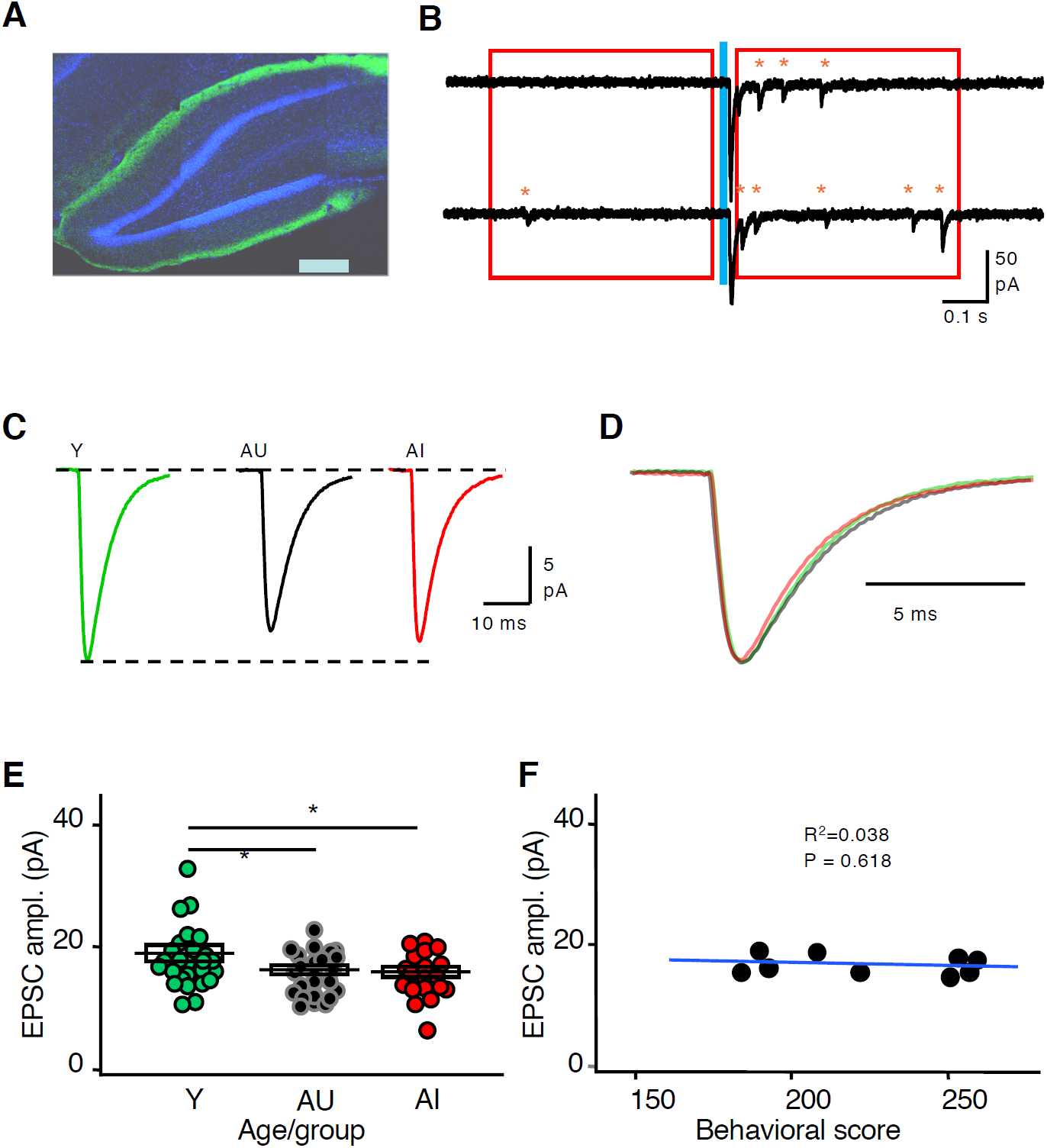
Aging reduces the postsynaptic strength of the inputs from lateral entorhinal cortex onto granule cells. **A)** Example of YFP expressing inputs (green) in the dentate gyrus of a rat injected with AAV-ChR2-EYFP in the lateral entorhinal cortex, granule cells were stained with DAPI (shown in green. Calibration: 300□m). **B)** The current traces are examples of optogenetically evoked (blue line) LEC→GC synaptic responses exhibiting Sr+2-induced desynchronization. Red boxes indicate the time windows used to detect isolated synaptic events (red asterisks) before and after the stimulation. **C**) The traces represent the computed quantal response for the LEC→GC averaged across all cells recorded in each of the three groups. The amplitude was larger in the young group (Y: green trace) than in the aged unimpaired (AU: black trace) or in the Aged impaired group (AI: red trace). **D)** The superimposed traces are the same one as in C but normalized to appreciate the similar kinetics. **E)** Quantal size amplitude of the LEC→GC mEPSCs for cells in the different age groups. Color code as in B. Boxes indicate group average ± SEM. **F)** In aged rats the average quantal size amplitude does not correlate with the behavioral score. The graph plots the mEPSC amplitude averaged by rat against the individual’s behavioral score. The blue line represents the best linear fit of the data.

Evoked excitatory events were quantified as described before (Petrus et al., 2014). A 400-ms window before LED was used for quantifying spontaneous desynchronized events (preLED), and a 400-ms window following a 50-ms delay from LED onset was used for quantifying LED-evoked desynchronized events (postLED). The average amplitude of the quantal events evoked (avQev) by the LED flash (520 nm, 3 msec whole field through the objective) was computed from the frequency (F) and amplitude (A) of isolated events recorded before (spontaneous events) and after stimulation (spontaneous+desynchronized evoked events) using the formula: avQev, = (Aafter x Fafter – Abef x Fbef)/ (Fafter –Fbef). The rate and amplitude of the quantal events were computed using the Mini Analysis Program (Synaptosoft) as previously described (Morales et al., 2002). For event discrimination we used an amplitude threshold of 3 times the RMS noise and a rise time<3 msec. The experimenter confirmed the events detected by the program.

To evaluate the E/I ratio, an internal pipette solution containing125 mM Cs-gluconate, 8 mM KCl, 1 mM EGTA, 10 mM HEPES, 4 mM ATP, 0.5 mM GTP; pH 7.4; 285-295 mOsm was used. Excitatory and inhibitory current reversal potentials were measured to be +10 mV and −55 mV, respectively, without compensating for the junction potential. A series of stimulations over a range of light intensities was delivered while holding at each reversal potential to generate an input-output curve for both EPSCs and IPSCs. Only the responses within the linear range of both I-O curves (stable E/I ratio) were used to calculate the evoked E/I ratio.

### Statistical analysis

Statistical significance was determined with Prism Graph Pad using ANOVA tests followed by Dunnet post hoc test, when the data was distributed normally (judged by the D’Agostino-Pearson normality test), or by the Kruskal-Wallis (K-W) test followed by the Dunn test.

All procedures were approved by the IACUC at Johns Hopkins University

## RESULTS

The aim of the study was to evaluate changes associated with cognitive aging in the synaptic inputs from the lateral entorhinal cortex (LEC) onto the granule cells (GC) in the dentate gyrus. To that end, we combined optogenetics, to specifically activate LEC input, with whole-cell methods to record excitatory and inhibitory postsynaptic currents (EPSC and IPSC, respectively). The measurements were done in slices prepared from young mature (6-month old) and aged (24-month old) outbred Long-Evans rats that had been behaviorally characterized in a standardized spatial learning protocol in a Morris water maze as described previously (see Methods), and subsequently infected with an AAV-ChR2 in the LEC approximately 6 weeks prior to the final experiment.

Figure 1 summarizes the behavioral assessment of the rats used in this study. Performance in the first trial (TT data point, Fig 1A), before the rats had experienced escaping to the hidden platform in the water maze, was similar in both age groups. Over the course of training, however, young rats were more proficient in learning to locate the platform. A two-way ANOVA (Age x Trial Block) showed that performance improved over the course of training (Trial Block, F(3,76) = 25.24, p = 0.001) but yielded a significant difference in overall performance between the two age groups (Age, F(1,76) = 10.94, p = 0.001). The interaction between Trial Block and Age was not significant (F(3,76) = 0.22, p = 0.882). The learning index scores (Fig 1B), computed from a key measure of search accuracy during interleaved probe trials, also differed according to age, with the young rats performing significantly better than the aged rats (Mann-Whitney U test = 28, p = 0.046). In agreement with previous research in this model, the aged rats displayed a range of outcomes, with a subset of aged rats performing on par with young adults and a substantial subset performing outside the range of young performance. Aged rats performing outside the range of the young group were designated as aged impaired (AI), while those performing on par with young adults (Y) were designated as aged unimpaired (AU). The cutoff used to identify AI and AU subgroups in the current study was consistent with normative data in this research population collected over many years.

### Age-related changes in response to input from LEC onto DG do not distinguish between AU and AI rats

The comparison of the strength of LEC→GC inputs between individuals using optogenetics in slices is limited by variations in the number of axons recruited, which in turn depends on contingencies like the extent of the viral transfection, the expression of ChR2 and the slice sectioning. Therefore, we focused the analysis on the quantification of intrinsic determinants of synaptic transmission, such as the quantal size of the postsynaptic response to the release of a single neurotransmitter vesicle. To that end, we used the divalent cation strontium (Sr^2+^) desynchronization approach that promotes the asynchronous release of neurotransmitters and allows individual quantal events (caused by the release of a single vesicle) to be resolved and quantified (asterisks in Fig 2B). Although it is not possible to determine whether a given isolated event is spontaneous or evoked, the average quantal size of the evoked responses in a given cell can be computed from the frequency and amplitude of isolated events recorded before (spontaneous events) and after stimulation (spontaneous + desynchronized evoked events. See Fig 2B and Methods for more details). The results, shown in Figure 2, indicate that age does not affect the shape of the quantal events evoked in granule cells. The computed events averaged across all cells in each of the three groups (Y, AU, AI) superimpose nicely when normalized for the amplitude (Fig 2D), and no significant differences were detected in the rise and decay times (see Table 1 for data and statistical analysis). The amplitude, on the other hand, was affected by age (Figs 2C and E). Compared to the young group (18.18 ± 0.96 pA), the average amplitude was significantly smaller in each of the aged subgroups, with no difference between the AU (15.49 ± 0.66 pA) and AI (15.47 ± 0.80 pA) subgroups of rats (Fig 2E; Tukey’s HSD post hoc tests with α set at 0.05). To further examine the difference from young adults across the spectrum of aged rat performance, we examined the relationship between the average event amplitude and the behavioral measure of performance among the aged rats. We averaged the event amplitude from all the cells recorded in a given aged individual and plotted this value against the individual’s behavioral performance in the water maze. Only individuals with three or more cells recorded were included in this analysis. As shown in Figure 2F, the individuals’ evoked event amplitude including all aged subjects did not correlate with behavioral performance, r^2^ = 0.038, p = 0.618. In sum, aging altered the event amplitude, yet in aged individuals, the average amplitude does not predict behavioral performance.

**Table 1.** Parameters of LEC-evoked quantal synaptic events.

| | N<br>rats/cells | Amplitude<br>(pA) | Rise time<br>(msec) | Decay tau<br>(msec) | RMS<br>(pA) | R input<br>(MΩ) | R access<br>(MΩ) |
| --- | --- | --- | --- | --- | --- | --- | --- |
| Y | 7/26 | 18.18±0.96 | 1.21±0.07 | 5.27±0.19 | 1.78±0.03 | 415±28 | 19.4±0.9 |
| AU | 7/27 | 15.49±0.66 | 1.36±0.07 | 5.45±0.11 | 1.81±0.03 | 571±41 | 20.9±0.9 |
| AI | 7/23 | 15.47±0.80 | 1.45±0.15 | 5.08±0.21 | 1.80±0.03 | 556±35 | 21.7±0.7 |
| Prob. |  | P=0.0262 | P=0.3185 | P=0.1572 | 0.9793 | 0.0049 | 0.0531 |
| Stats. |  | F (2,72)<br>3.833 | KW<br>2.29 | KW<br>3.7 | KW<br>0.042 | KW<br>10.62 | KW<br>5.87 |

In a subsequent set of studies, we evaluated possible age-related alterations in presynaptic release using paired-pulse stimulation, which provides an approximate estimate of changes in release probability. In these experiments synaptic responses were evoked with paired light pulses at interstimulus intervals (isi) of 50ms, 100 ms and 200ms (Fig 3A). The results indicate that aging altered the paired-pulse amplitude ratio (PPR: response2/response1) of the responses, but only those evoked with a 100 ms isi. A two-way ANOVA confirmed the differences in time intervals, F(2,162) = 37.55, p = 0.001 among the three groups, F(2,81) = 5.847, p = 0.004. Although no significant interaction between group and time interval was detected, F(4, 162) = 1.811, p = 0.129, Tukey’s post hoc tests (with α set at 0.05) indicated that at the 100ms isi, the average PPR in each of the aged groups (AU, AI) was smaller than the young group (Y), with no difference between the two aged subgroups. As in the case of the event amplitudes, the average value of the PPR at 100 ms isi per rat did not correlate with the individual’s behavioral score (Fig 3C; r^2^ = 0.084, p = 0.447). The smaller PPR in the aged groups is consistent with an increased presynaptic release probability, which could serve as a compensation to maintain input strength in the face of reduced postsynaptic responsiveness. In any case, it must be noted that although age affects both pre- and postsynaptic measures of synaptic efficacy, none of these changes distinguished AI from AU individuals.

**Figure 3.**
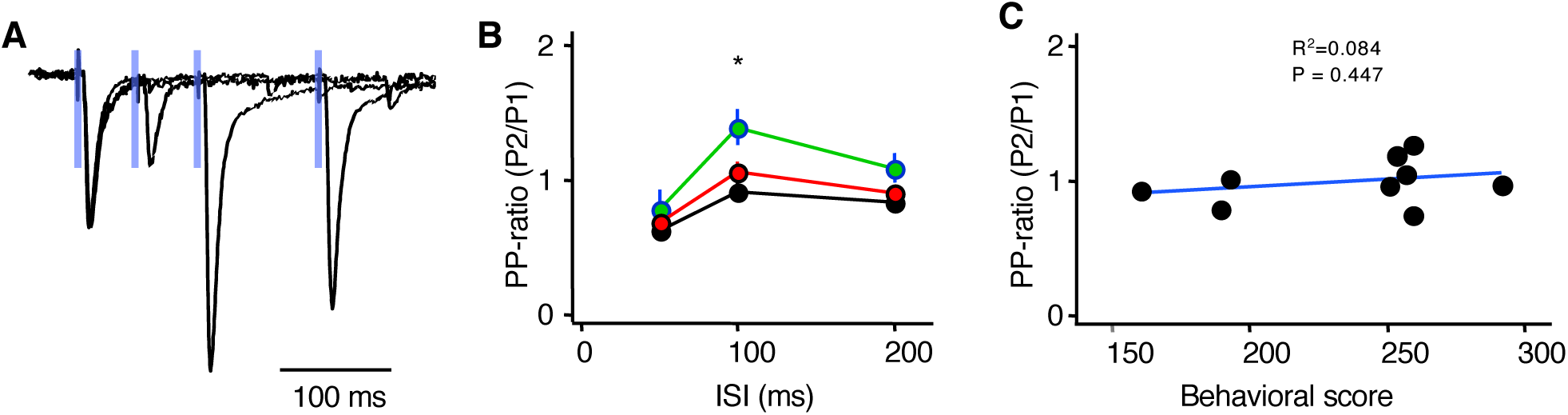
Aging reduces the paired-pulse facilitation of the inputs from lateral entorhinal cortex onto granule cells. **A)** Superimposed traces are examples of optogenetically evoked responses to paired pulse stimulation of 3 different intervals (50ms, 100ms, 200 ms). Blue line indicates the 3 ms light pulse. **B)** Paired pulse response ratio (PPR = Response2/Response1) is increased in the Y group compared to the two aged groups. **C)** The paired-pulse ratio obtained at 100 ms interval does not correlate with the individual’s behavioral score. The graph plots the PPR averaged by rat against the individual’s behavioral score. The blue line represents the best linear fit of the data.

### Increased Excitation/Inhibition Ratio in the AI rats

The balance of synaptic excitation and inhibition is crucial for neural function in general, and for pattern separation in the DG in particular. Granule cells in the DG are subjected to strong feedforward inhibition recruited disynaptically by entorhinal inputs. Therefore, we asked whether age affects the ratio of synaptic excitation and inhibition (E/I ratio) evoked by optogenetic activation of LEC inputs. To that end, we recorded evoked EPSCs and IPSCs in the same granule cell by holding the membrane at the reversal potential for GABA (−50mV) and AMPA receptors (0mV), respectively. As illustrated in Figure 4A, substantial disynaptic feedforward inhibition occurs following the EPSC. Since the E/I ratio varies with stimulation intensity (Hsu-Lien 2016; Morales et al., 2002), we stimulated each cell at a range of intensities to determine the intensities over which the E/I ratio is stable (Fig 4B). The findings indicate that age does affect the E/I ratio of the responses evoked by LEC inputs (Kruskall-Wallis = 16.77, p = 0.001), with a change occurring specifically in the AI group, which exhibited the largest values for the E/I ratio (AI: 0.583 ± 0.068, n = 23; AU: 0.251 ± 0.038; n = 20; Y: 0.307 ± 0.037; n = 21). A Dunn’s test (with α set at 0.05) confirmed the significance of the differences between AI and each of the other groups. Moreover, analysis of the relationship between the E/I ratio and behavioral performance among the aged rats was significantly correlated (Fig 4C; r^2^ = 0.601, p = 0.042). These results are consistent with the notion that the integrity of inhibitory circuits is crucial for feedforward function in the DG.

**Figure 4.**
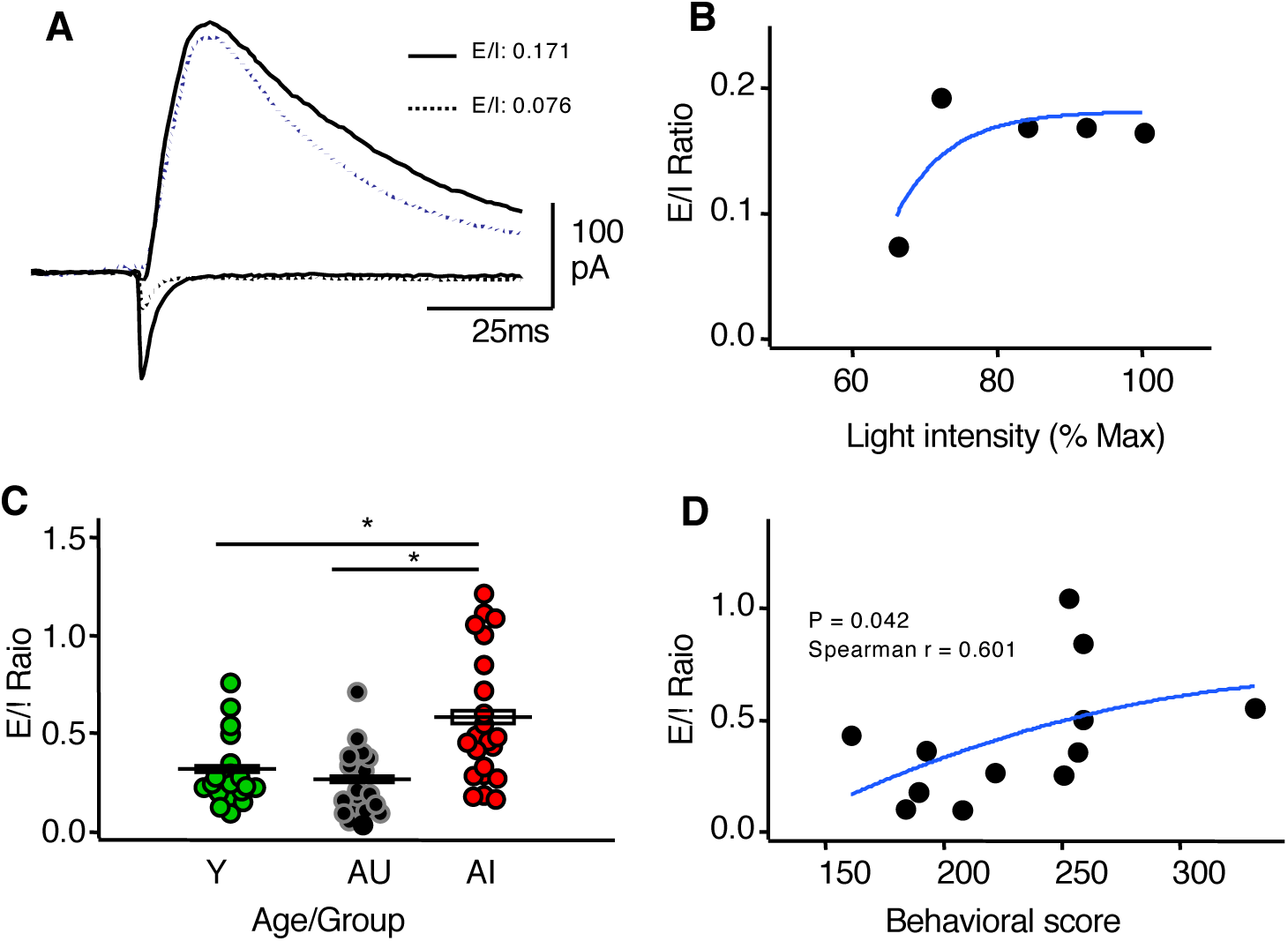
Larger excitation/inhibition ratio of the LEC→GC inputs in aged impaired rats. **A)** The traces are examples of excitatory synaptic responses (at a holding potential of −55 mV: lower traces) and inhibitory responses (recorded at +10 mV: upper traces) and evoked by light stimulus at lower (dotted line) and higher (solid line) intensity values. Note the different excitatory/ inhibitory (E/I) ratio obtained at the two intensities. **B)** Example showing that the E/I ratio asymptotes at increasing stimulus intensity. **C)** The E/I ratio is larger in the AI group compared to the AIU and Y groups. Boxes indicate group average ± SEM. **D)** The graph plots the individual’s behavioral score versus the average E/I ratio of the cells recorded in that individual. The two variables correlate as assessed by the Spearman test. The blue lines in B and D are drawn for visual purposes.

## DISCUSSION

The current findings indicate that aging alters the properties of excitatory LEC inputs onto granule cells. Some of those findings did not distinguish the cognitive characterization for individual differences among the aged rats. Indeed, such alterations appeared to be generally related to aging and unlikely to serve as critical features accounting for individual differences in hippocampal-dependent function. On the other hand, a profile in E/I balance in response to LEC stimulation distinguished the AI rats from both Young and AU cohorts. This finding adds to other AI effects that distinguish aging with impairment from young adults and aged rats with preserved cognitive function in this model (e.g. Smith et al. 2000; Lee et al 2005; Boric et al. 2007; Stranahan et al. 2010; Yang et al. 2013; Spiegel et al. 2013; Haberman et al. 2017).

An opposing effect on pre and postsynaptic strength could reflect the engagement of compensatory mechanisms to maintain overall synaptic strength. Since those synaptic changes do not correlate with differences in behavioral performance that distinguish AI and AU rats their contribution would not appear to be critical in differentiating cognitive outcomes We believe this is a significant negative finding because changes in the amplitude of miniature synaptic responses have often been implicitly implicated in behavioral deficits observed in models of aging and neurodegenerative diseases (Chang et al., 2006; Gocel and Larson, 2013). The lack of correlation with performance in the LEC-evoked miniature events reported here suggests that is not always the case and emphasizes the importance of behavioral corroborations especially in studies of circuitry where differences have been found to be tightly associated with cognitive status.

In contrast to the absence of correlation between measures of excitatory synaptic strength of the LEC inputs and behavioral performance in the aged rats, the relative strength of synaptic inhibition recruited by LEC activation was markedly reduced in the AI rats. Reduced feedforward synaptic inhibition relative to excitation dovetails well with multiple observations of increased activation and hyperactivity in the DG/CA3 region of cognitive impaired aged individuals, both in rodents and humans (Wilson et al., 2004; Yassa et al., 2010; Bakker et al., 2012; Leal and Yassa, 2015). These findings also fit with the notion that an increase in net activation could limit the capacity to support sparse coding in the DG, considered an essential condition for pattern separation in the memory encoding process.

The exact mechanisms underlying the reduced feedforward inhibition remain to be determined. The primary inhibitory neurons recruited by LEC inputs are fast spiking parvalbumin positive interneurons (PV-IN) (Hsu et al., 2016), and the molecular layer perforant pathway (MOPP) cells, that target the dendrites of GC (Li et al., 2013). The fast rise time of the inhibitory response reported here (less than 5 ms rise time) is consistent with a substantial recruitment of PV-INS, providing perisomatic inhibition, which occurs faster than the MOPP cells targeting distal dendrites (Li et al., 2013). An attractive and simple possibility then, is that the recruitment of PV-INs by LEC inputs is reduced in the AI rats. Other scenarios are certainly plausible, for example impaired interneuron excitability. A reduced recruitment of PV-INs, however, would be consistent with reports implicating dysfunctional PV-IN circuitry in aging and the early stages of Alzheimer’s disease in rodent models (Verret et al., 2012; Iaccarino et al., 2016; Kann, 2016), and humans (Xiao et al., 2017).

The reduced recruitment of feedforward inhibition in AI individuals complements previous reports that alterations in hilar cells providing feedback inhibition onto GC neurons are also associated with impaired performance in aging. Experimental reduction of SOM HIPP interneurons has been demonstrated to play a critical role in DG function by limiting granule cell activation. With respect to the current findings, those somatostatin positive HIPP interneurons are seemingly not recruited in a disynaptic fashion by entorhinal inputs (Hsu et al., 2016) but also undergo an age-dependent loss associated with memory impairment ((Andrews-Zwilling et al. 2010; 2012; Koh et al. 2014), including an AI selective loss in the current model of individual differences in outbred Long Evans rats (Spiegel et al.2013). Thus, aging appears to challenge key mechanisms that contribute to the proficient encoding of episodic memory in the circuitry of the DG.

Studies of individual differences in neurocognitive aging, such as the current investigation, demonstrate that such neurobiological changes, however, are not an invariable outcome of aging. In contrast with the individuals that are behaviorally impaired, subjects with preserved cognitive performance have a maintenance in the overall functional integrity of the LEC→GC network. Indeed, other studies using this model of neurocognitive aging have shown that AU rats have enhanced postsynaptic synaptic inhibition in granule cells and enhanced tonic inhibition in CA1 pyramidal cells (Tran et al., 2018). Moreover, preserved cognitive performance in this study population is associated with increased task-induced expression of gene markers of synaptic inhibition across hippocampal subfields in AU rats compared to young adults (Branch et al., 2019).

In sum, our analysis of the LEC inputs to granule cells indicates a reduced recruitment of synaptic inhibition, rather than changes in direct synaptic excitation correlate with reduced performance in aging. Indeed, the current investigation adds to a growing body of evidence that the functionality of inhibitory circuits may bidirectionally contribute to individual differences in cognitive outcomes in aging. A loss in such function is associated with impairment while recruitment mediated via inhibitory mechanisms may contribute to resilience against functional loss in the aged brain.

## ACKNOWLEDGMENTS

This work was supported by National Institute on Aging/National Institutes of Health Grants AG009973-22 to MG and AG009973-24 to AK. MB was supported by NIH T32HL110952.

## REFERENCES

Andrews-Zwilling Y, Bien-Ly N, Xu Q, Li G, Bernardo A, Yoon SY, Zwilling D, Yan TX, Chen L, Huang Y (2010) Apolipoprotein E4 causes age-and Tau-dependent impairment of GABAergic interneurons, leading to learning and memory deficits in mice. Journal of Neuroscience, 30(41), pp. 13707–13717.

Andrews-Zwilling Y, Gillespie AK, Kravitz AV, Nelson AB, Devidze N, Lo I, Yoon SY, Bien-Ly N, Ring K, Zwilling D, and Potter GB (2012) Hilar GABAergic interneuron activity controls spatial learning and memory retrieval. PloS one, 7(7), p. e40555.

Bakker A, Krauss GL, Albert MS, Speck CL, Jones LR, Stark CE, Yassa MA, Bassett SS, Shelton AL, Gallagher M (2012) Reduction of hippocampal hyperactivity improves cognition in amnestic mild cognitive impairment. Neuron 74:467–474.

Bondareff W, Geinisman Y(1976). Loss of synapses in the dentate gyrus of the senescent rat. American journal of anatomy, 145(1), 129–136.

Boric K, Muñoz P, Gallagher M, Kirkwood A. Potential adaptive function for altered long-term potentiation mechanisms in aging hippocampus. J Neurosci. 28:8034–8039.

Branch A, Monasterio A, Blair G, Knierim JJ, Gallagher M, Haberman RP (2019) Aged rats with preserved memory dynamically recruit hippocampal inhibition in a local/global cue mismatch environment. Neurobiol Aging 76:151–161.

Chang EH, Savage MJ, Flood DG, Thomas JM, Levy RB, Mahadomrongkul V, Shirao T, Aoki C, Huerta PT (2006) AMPA receptor downscaling at the onset of Alzheimer’s disease pathology in double knockin mice. Proc Natl Acad Sci U S A 103:3410–3415.

Foster TC, Barnes CA, Rao G, McNaughton BL (1991) Increase in perforant path quantal size in aged F-344 rats. Neurobiol Aging 12:441–448.

Gallagher M (1993) Issues in the development of models for cognitive aging across primate and nonprimate species. Neurobiol Aging 14:631–633.

Gallagher M, Colantuoni C, Eichenbaum H, Haberman RP, Rapp PR, Tanila H, Wilson I A (2006). Individual differences in neurocognitive aging of the medial temporal lobe. Age, 28(3), 221–233.

Gallagher M, Burwell R, Burchinal M (2015) Severity of spatial learning impairment in aging: development of a learning index for performance in the Morris water maze.

Gocel J, Larson J (2013) Evidence for loss of synaptic AMPA receptors in anterior piriform cortex of aged mice. Front Aging Neurosci 5:39.

Haberman RP, Koh MT, Gallagher M (2017) Heightened cortical excitability in aged rodents with memory impairment. Neurobiology of aging, 54, 144–151.

Haberman RP, Branch A, Gallagher M (2017) Targeting neural hyperactivity as a treatment to stem progression of late-onset Alzheimer’s disease. Neurotherapeutics, 14(3), 662–676.

Hsu TT, Lee CT, Tai MH, Lien CC (2016) Differential Recruitment of Dentate Gyrus Interneuron Types by Commissural Versus Perforant Pathways. Cereb Cortex 26:2715–2727.

Iaccarino HF, Singer AC, Martorell AJ, Rudenko A, Gao F, Gillingham TZ, Mathys H, Seo J, Kritskiy O, Abdurrob F, Adaikkan C, Canter RG, Rueda R, Brown EN, Boyden ES, Tsai LH (2016) Gamma frequency entrainment attenuates amyloid load and modifies microglia. Nature 540:230–235.

Kann O (2016) The interneuron energy hypothesis: Implications for brain disease. Neurobiol Dis 90:75–85.

Khan U, Liu L, Provenzano F, Berman D, Profaci C, Sloan R, Mayeux R, Duff K, Small S (2014) Molecular drivers and cortical spread of lateral entorhinal cortex dysfunction in preclinical Alzheimer’s disease. Nat Neurosci in press.

Koh MT, Spiegel AM, Gallagher M (2014). Age-associated changes in hippocampal-dependent cognition in D iversity O utbred mice. Hippocampus, 24(11), 1300–1307.

Kulason S, Tward DJ, Brown T, Sicat CS, Liu CF, Ratnanather JT, Younes L, Bakker A, Gallagher M, Albert M, Miller MI (2019) Cortical thickness atrophy in the transentorhinal cortex in mild cognitive impairment. NeuroImage: Clinical, 21, p.101617.

Lee HK, Min SS, Gallagher M, Kirkwood A. (2005). NMDA receptor-independent long-term depression correlates with successful aging in rats. Nat Neurosci. 8:1657–1659.

Leal SL, Yassa MA (2015) Neurocognitive Aging and the Hippocampus across Species. Trends Neurosci 38:800–812.

Leutgeb JK, Leutgeb S, Moser MB, Moser EI (2007) Pattern separation in the dentate gyrus and CA3 of the hippocampus. Science 315:961–966.

Li Y, Stam FJ, Aimone JB, Goulding M, Callaway EM, Gage FH (2013) Molecular layer perforant path-associated cells contribute to feed-forward inhibition in the adult dentate gyrus. Proc Natl Acad Sci U S A 110:9106–9111.

McHugh TJ, Jones MW, Quinn JJ, Balthasar N, Coppari R, Elmquist JK, Lowell BB, Fanselow MS, Wilson MA, Tonegawa S (2007) Dentate gyrus NMDA receptors mediate rapid pattern separation in the hippocampal network. Science 317:94–99.

Morales B, Choi SY, Kirkwood A (2002) Dark rearing alters the development of GABAergic transmission in visual cortex. J Neurosci 22:8084–8090.

Neunuebel JP, Knierim JJ (2014) CA3 retrieves coherent representations from degraded input: direct evidence for CA3 pattern completion and dentate gyrus pattern separation. Neuron 81:416–427.

Palop JJ, Chin J, Roberson ED, Wang J, Thwin MT, Bien-Ly N, Yoo J, Ho KO, Yu GQ, Kreitzer A, Finkbeiner S, Noebels JL, Mucke L (2007) Aberrant excitatory neuronal activity and compensatory remodeling of inhibitory hippocampal circuits in mouse models of Alzheimer’s disease. Neuron 55:697–711.

Petrus E, Isaiah A, Jones AP, Li D, Wang H, Lee HK, Kanold PO. (2014) Crossmodal induction of thalamocortical potentiation leads to enhanced information processing in the auditory cortex. Neuron 81: 664–673.

Rapp PR, Gallagher M (1996) Preserved neuron number in the hippocampus of aged rats with spatial learning deficits. Proc Natl Acad Sci U S A 93:9926–9930.

Rapp PR, Stack EC, Gallagher M (1999) Morphometric studies of the aged hippocampus: I. Volumetric analysis in behaviorally characterized rats. J Comp Neurol 403:459–470.

Scheff SW, Price DA, Schmitt FA, Mufson EJ (2006). Hippocampal synaptic loss in early Alzheimer’s disease and mild cognitive impairment. Neurobiology of aging, 27(10), 1372–1384.

Smith TD, Adams MM, Gallagher M, Morrison JH, Rapp PR (2000) Circuit-specific alterations in hippocampal synaptophysin immunoreactivity predict spatial learning impairment in aged rats. Journal of Neuroscience, 20(17), 6587–6593.

Spiegel AM, Koh MT, Vogt NM, Rapp PR, Gallagher M (2013). Hilar interneuron vulnerability distinguishes aged rats with memory impairment. Journal of Comparative Neurology, 521(15), 3508–3523.

Stranahan AM, Mattson MP (2010) Selective vulnerability of neurons in layer II of the entorhinal cortex during aging and Alzheimer’s disease. Neural Plast 2010:108190.

Stranahan AM, Haberman RP, Gallagher M (2010) Cognitive decline is associated with reduced reelin expression in the entorhinal cortex of aged rats. Cereb Cortex 21:392–400.

Stranahan AM, Salas-Vega S, Jiam NT, Gallagher M (2011) Interference with reelin signaling in the lateral entorhinal cortex impairs spatial memory. Neurobiol Learn Mem 96:150–155.

Tran T, Gallagher M, Kirkwood A (2018) Enhanced postsynaptic inhibitory strength in hippocampal principal cells in high-performing aged rats. Neurobiol Aging 70:92–101.

Tward DJ, Sicat CS, Brown T, Bakker A, Gallagher M, Albert M, Miller M and Alzheimer’s Disease Neuroimaging Initiative (2017) Entorhinal and transentorhinal atrophy in mild cognitive impairment using longitudinal diffeomorphometry. Alzheimer’s & Dementia: Diagnosis, Assessment & Disease Monitoring, 9, pp.41–50.

Verret L, Mann EO, Hang GB, Barth AM, Cobos I, Ho K, Devidze N, Masliah E, Kreitzer AC, Mody I, Mucke L, Palop JJ (2012) Inhibitory interneuron deficit links altered network activity and cognitive dysfunction in Alzheimer model. Cell 149:708–721.

Wilson IA, Ikonen S, Gureviciene I, McMahan RW, Gallagher M, Eichenbaum H, Tanila H (2004) Cognitive aging and the hippocampus: how old rats represent new environments. J Neurosci 24:3870–3878.

Wilson IA, Ikonen S, Gurevicius K, McMahan RW, Gallagher M, Eichenbaum H, Tanila H (2005) Place cells of aged rats in two visually identical compartments. Neurobiol Aging 26:1099–1106.

Wilson IA, Ikonen S, Gallagher M, Eichenbaum H, Tanila H (2005). Age-associated alterations of hippocampal place cells are subregion specific. Journal of Neuroscience, 25(29), 6877–6886.

Xiao MF, Xu D, Craig MT, Pelkey KA, Chien CC, Shi Y, Zhang J, Resnick S, Pletnikova O, Salmon D, Brewer J, Edland S, Wegiel J, Tycko B, Savonenko A, Reeves RH, Troncoso JC, McBain CJ, Galasko D, Worley PF (2017) NPTX2 and cognitive dysfunction in Alzheimer’s Disease. Elife 6.

Yang S, Megill A, Ardiles AO, Ransom S, Tran T, Koh MT, Lee HK, Gallagher M, Kirkwood A. (2013). Integrity of mGluR-LTD in the associative/commissural inputs to CA3 correlates with successful aging in rats. J Neurosci. 33:12670–12678.

Yassa MA, Stark SM, Bakker A, Albert MS, Gallagher M, Stark CE (2010) High-resolution structural and functional MRI of hippocampal CA3 and dentate gyrus in patients with amnestic Mild Cognitive Impairment. Neuroimage 51:1242–1252.

Yassa MA, Muftuler LT, Stark CE (2010). Ultrahigh-resolution microstructural diffusion tensor imaging reveals perforant path degradation in aged humans in vivo. Proceedings of the National Academy of Sciences, 107(28), 12687–12691.

Yassa MA, Lacy JW, Stark SM, Albert MS, Gallagher M, Stark CE (2011) Pattern separation deficits associated with increased hippocampal CA3 and dentate gyrus activity in nondemented older adults. Hippocampus 21:968–979.

